# Non-invasive prenatal testing as a valuable source of population specific allelic frequencies

**DOI:** 10.1101/348466

**Authors:** J Budis, J Gazdarica, J Radvanszky, M Harsanyova, I Gazdaricova, L Strieskova, R Frno, F Duris, G Minarik, M Sekelska, T Szemes

## Abstract

Low-coverage massively parallel genome sequencing for non-invasive prenatal testing (NIPT) of common aneuploidies is one of the most rapidly adopted and relatively low-cost DNA tests. Since aggregation of reads from a large number of samples allows overcoming the problems of extremely low coverage of individual samples, we describe the possible re-use of the data generated during NIPT testing for genome scale population specific frequency determination of small DNA variants, requiring no additional costs except of those for the NIPT test itself. We applied our method to a data set comprising of 1,548 original NIPT test results and evaluated the findings on different levels, from *in silico* population frequency comparisons up to wet lab validation analyses using a gold-standard method. The revealed high reliability of variant calling and allelic frequency determinations suggest that these NIPT data could serve as valuable alternatives to large scale population studies even for smaller countries around the world.

## Introduction

Although the costs of sequencing are continually dropping (Erlich 2015), large-scale human genome-related projects (Carrasco-Ramiro et al. 2017) still remain to be associated with substantial costs and a certain timeframe to complete. Further data aggregation efforts are, however, still in place to increase resolution and improve power at low allele frequencies (Lek et al. 2016). On the other hand, a low-cost genomic test readily used in routine clinical practice for determination of common fetal chromosomal aberrations and selected copy-number variants is becoming commonplace (Minear et al. 2015; Gregg et al. 2016). Non-invasive prenatal testing (NIPT) most commonly uses very low-coverage massively parallel whole-genome sequencing of total plasma DNA of pregnant women (Minarik et al. 2015). Although high-quality single nucleotide variant (SNVs) and small insertion-deletion (indels) calls can be observed even in individual reads, reliable genotyping through one mapped read per genomic position cannot be considered appropriate in individual patients. Hypothetically, however, the vast amount of data generated during NIPT testing worldwide (Shendure et al. 2017) could be used for whole-genome-scale population specific frequency determination of small sequence variants. The rationale behind our vision lies in overcoming the problems of extremely low coverage of individual samples by aggregation of reads from a large cohort routinely tested. To evaluate the possibilities/limitations of such an approach, we analysed data generated by massively parallel low-coverage whole-genome sequencing of plasma DNA of 1,548 pregnant women undergoing NIPT procedure in Slovakia.

## Results

To evaluate the above mentioned possibilities, 1,548 individual binary alignment map (BAM) files were analysed, revealing a median per sample genome coverage of 20.36% (0.2x), while 16.99% was represented by single covered positions in individual samples (Fig. 1, 2 and 3a).

**Figure 1:**
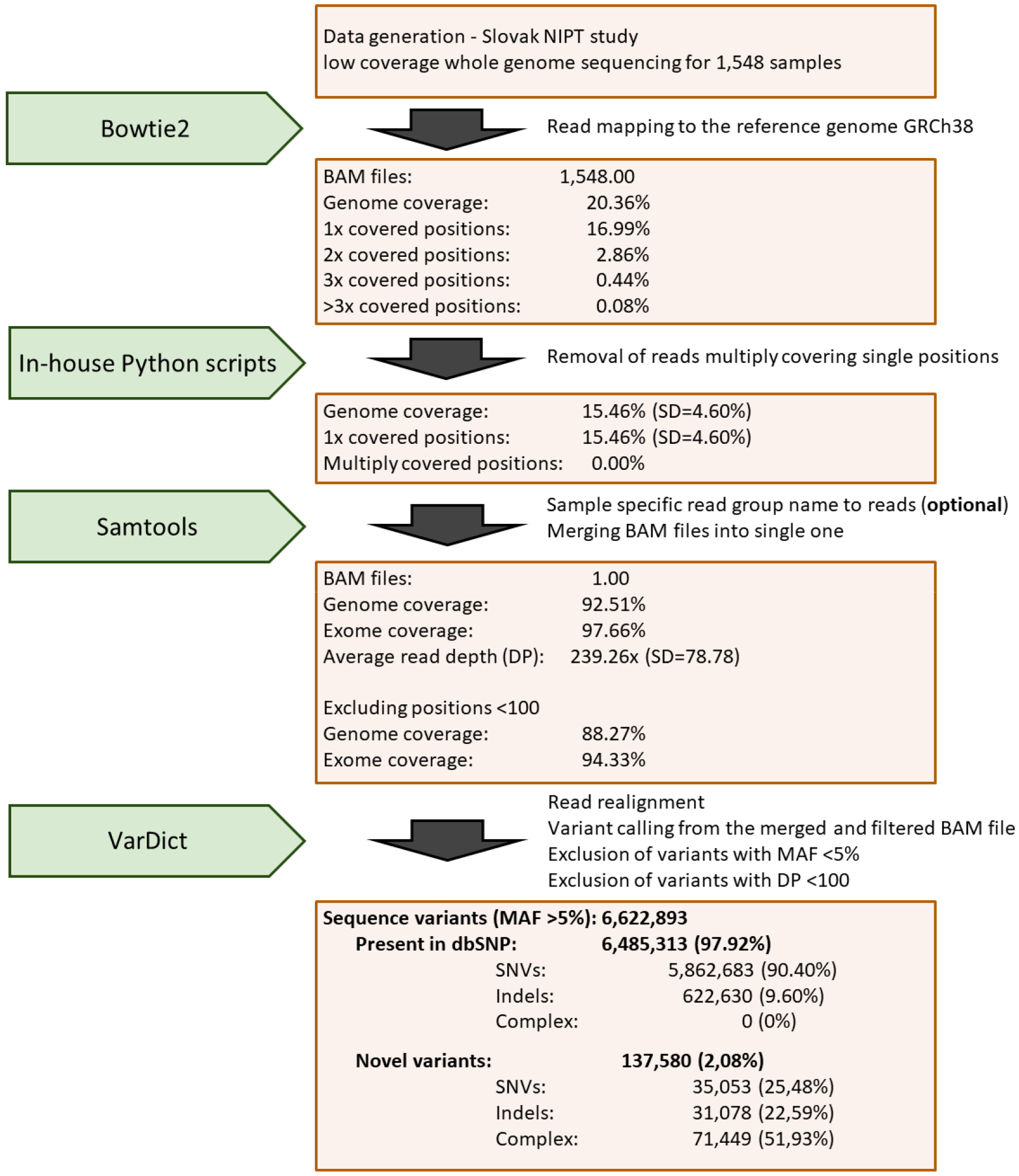
Schematic sequential representation of the performed analytical steps, the applied tools, filters and statistical results. BAM = binary alignment map; DP = depth of coverage; GRCh = Genome Reference Consortium human; MAF = minor allele frequency; NIPT = non-invasive prenatal testing; SD = standard deviation; SNVs = single nucleotide variants.

**Figure 2:**
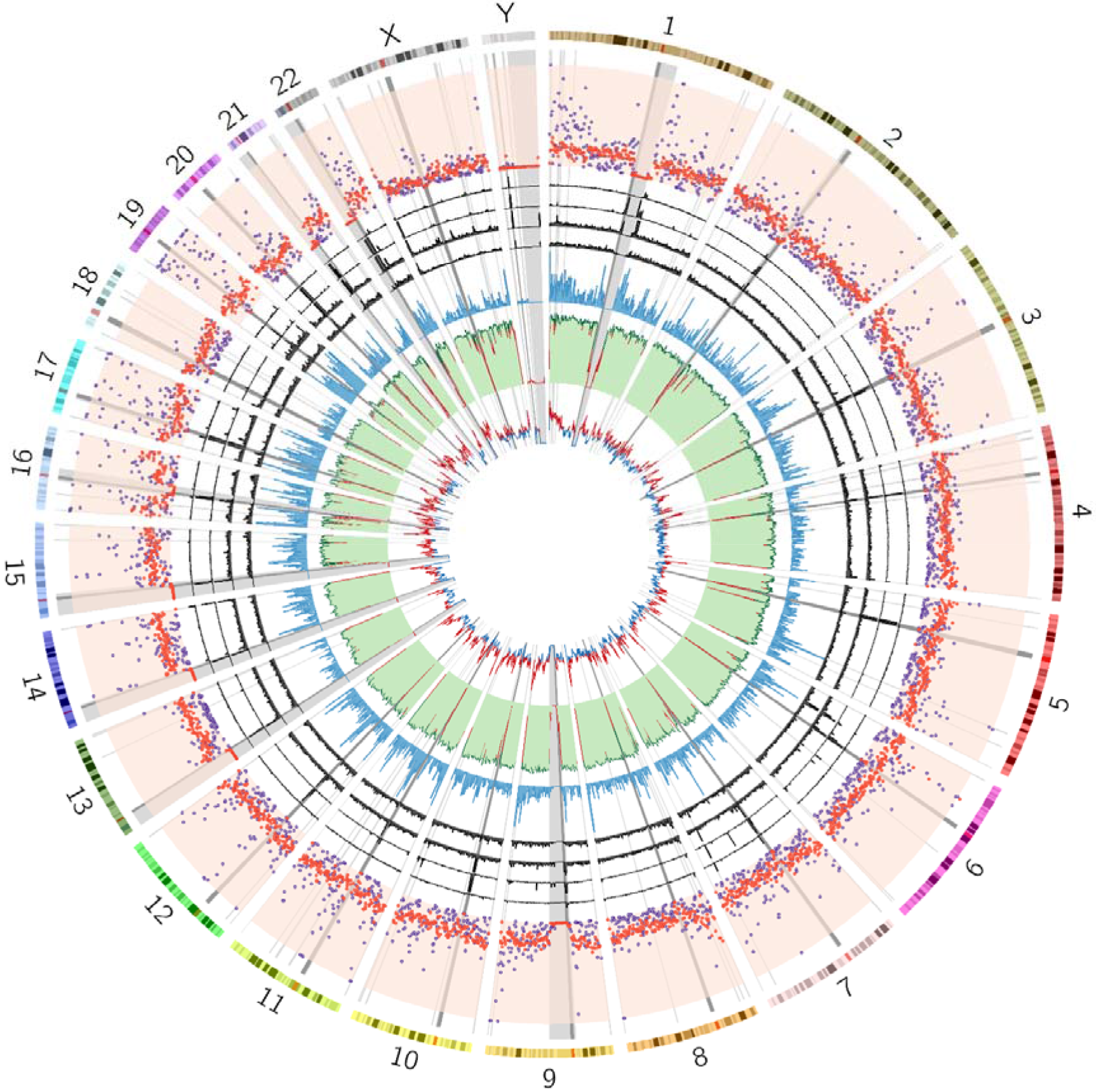
Characteristics of genome coverage and variant distributions, clustered by 1,000,000 bases. Dark grey regions = centromeres; light grey regions = unmappable genomic regions which are not assembled in the reference genome (N regions). Track numbering from the inner circle (axis of each track corresponds from low value to high value from the inside to the outside): **Track 1:** GC content of the genome sequence; **Track 2:** Genome coverage and coverage depth. Distribution of the uncovered regions strongly correlates with the unassembled regions of the genome consisting mainly of telomeres, centromeres, short-arms of acrocentric chromosomes (chr13, 14, 15, 21, 22, Y) and large heterochromatic regions of chr1, 9, 16, Y (Li and Freudenberg 2014); **Track 3:** Ensembl-based gene density of the genome. **Track 4-7:** Novel variant positions, lacking records in dbSNP (137,580), separated to complex variants (**Track 4**), variants in repetitive regions (**Track 5**), variants in regions that are not present in GRCh37 (**Track 6**) and novel variants with so far unidentified aetiology marked as “unresolved” (**Track 7**); **Track 8:** Variant density for those 6,485,313 variants that have frequency higher than 5% and are already present in dbSNP (purple dots = ExAC data set; red dots = our data set).

**Figure 3:**
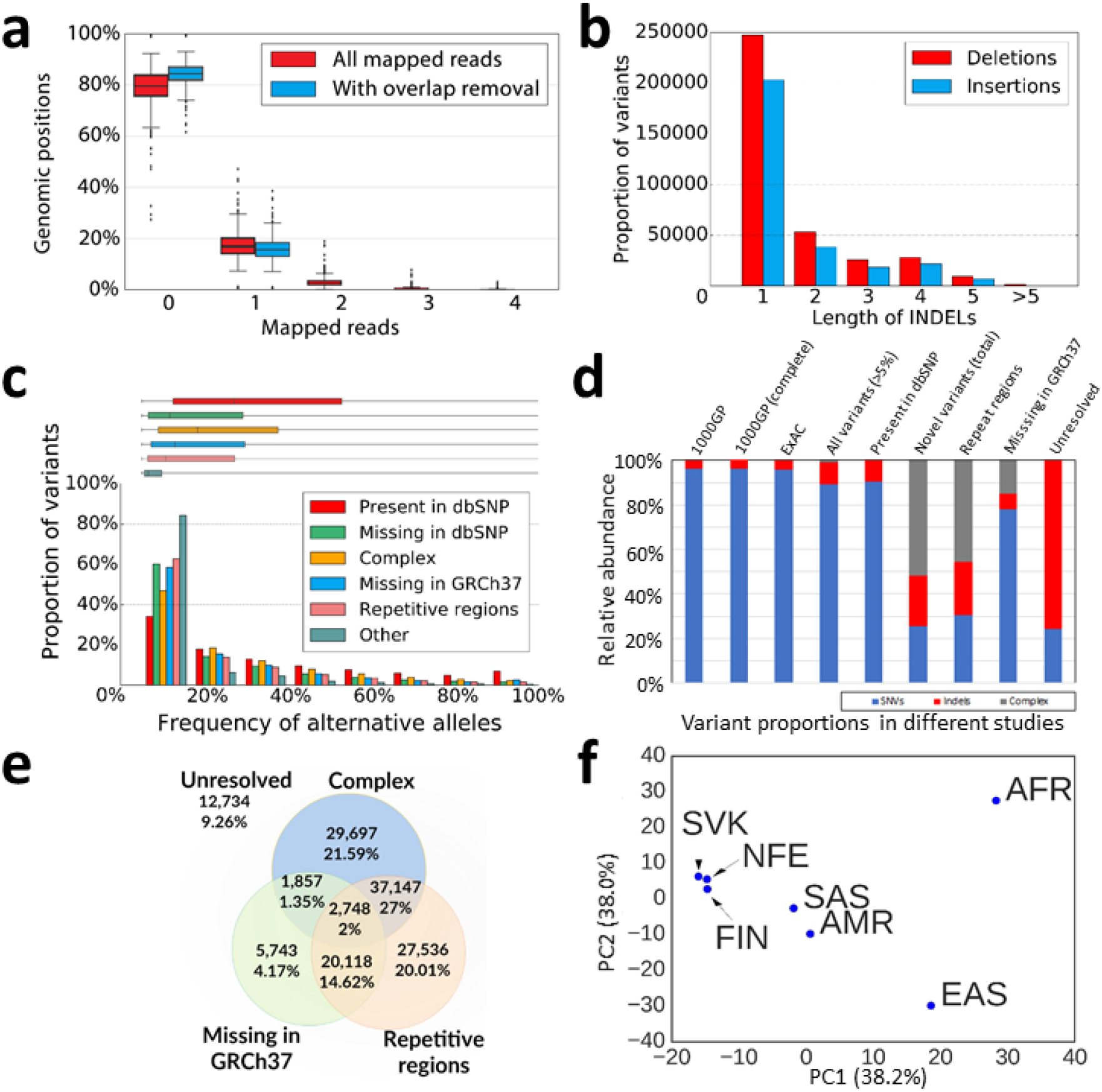
Graphical representation of statistical results. (**a**) Read coverage distribution of 1548 sequenced samples. We show the portions of the genome with the number of aligned reads from zero to four. Red boxes show samples before removal of the overlapping reads, while the blue ones represent filtered samples whose overlapped reads were removed. (**b**) Frequency distributions of insertions and deletions in our variant set ordered by size. (**c**) Graphical comparison of alternative allele frequencies identified in our sample. (**d**) Relative proportion of variant types found in previous studies (1000 Genomes Project and ExAC) as well as in our data set for all variants, those present in dbSNP and those not identified in dbSNP (and the three main categories of these “novel” variants). Indel variants in the 1000 Genomes project data set based only on dinucleotide variants (Altshuler et al. 2012; Auton et al. 2015). Moreover, neither 1000GP nor ExAC mentioned complex variant types. (**e**) Relative portion of “novel” variant subgroups with their overlaps. (**f**) Principal component analysis (PCA) for the comparison of allelic frequencies (both SNVs and indels) in our sample set and six different ExAC populations. Based on 71,235 variants simultaneously identified in each data subset with MAF higher than 5%. AFR = African/African American, AMR = American (Latino), EAS = East Asian, FIN = Finnish, NFE = Non-Finnish European, SAS = South Asian, SVK = Slovak.

Multiply covered regions in a single individual, however, pose a problem in determination of the total number of alleles for frequency calculations, thus having a potential to skew the resulting calculated allelic frequencies for variants detected in these particular regions. Since our specific simulations, performed to estimate the effect of this factor, suggested substantial increase of simulated MAF variance caused by duplicates (Suppl. Fig. 1), we decided to remove all overlapping reads to ensure uniqueness of observed alleles. Reads from individual BAM files were filtered in such way that only the first of overlapping reads were kept for further analysis. Due the fact that one whole read was removed in each pair, this step led also to removal of not only the overlapped parts but also some uniquely mapped positions. This was the reason why 16.99% of uniquely mapped positions came down to 15.46% following this step. Optional modification of the BAM files consisted of labelling of each individual BAM file with sample-specific read group name allowing both merging BAM files without any irreversible loss of specific information and also later verification analyses. Merging of individual BAM files into one “master BAM” led to a 92.51% genome coverage. Although, 4.24% of the covered genomic positions were found to have read depth below our arbitrarily set threshold of 100 mapped reads and were excluded from further analyses (Suppl. Fig. 2). Finally, altogether 88.27% of the reference genome was covered with sufficient read depth for variant calling.

Following variant calling from the merged BAM file, because of statistical reasons, we decided to further analyse only variants with detected minor allele frequencies (MAF) above 5%. Under these settings we identified 6,622,893 (0.21% of the genome) unique genomic positions with potential sequence variants detected. From these, altogether 6,485,313 (97.92%) variants were already known and described in dbSNP, while 137,580 (2.08%) of them were found to be novel variants, not present in dbSNP. In general, identified variants included both SNVs, complex variants and indels, while the latter group consisted of variants having lengths <6bp, with typically shorter indels being the most common ones (Fig. 3b).

The first step of our subsequent verification procedure included principal component analyses based comparisons of allelic frequency distributions, identified in our population sample (Slovak population located in Central Europe) to the six ExAC populations that placed our sample set most closely to the two European ExAC populations, i.e. to the Finnish and non-Finnish European population sample sets (Fig. 3f; Suppl. Fig. 3, 4). Next, we compared also population frequencies of variants in a single particular gene (chloride voltage-gated channel 1; *CLCN1*; UniProtKB_P35523) that further suggested high reliability of variant calling and frequency determination. Specifically, we identified all but five of ExAC *CLCN1* variants, having frequency >5%, in our data set too, with calculated population frequencies highly similar to those from ExAC. The exceptions were found to have ExAC frequencies around 5% (depending on population), while three of these we identified in our original data set too, although were filtered out due slightly lower than 5% frequency (Suppl. Tab. 1). In addition to these *in silico* verification approaches, we validated 87 positions in five polymorphic genomic loci of 58 randomly selected samples of our sample set, from which genomic DNA was available to validation purposes. These validation analyses revealed Sanger determined genotypes fully compatible with the NIPT derived allele for each of the particular loci (Suppl. Tab. 2).

## Discussion

Although NIPT is generally performed using whole-genome sequencing with genome coverage well below 1x, multiply covered regions can generally be identified in individual samples. Since these cause problems in determination of the total number of alleles for frequency calculations, we filtered out overlapping reads from our individual data sets that, as anticipated, in turn led to a lower total genome coverage. Although individual BAM files suffered by this filtration, in terms of overall genome coverage, after this filtering step we were able to set the total allele count as one allele per individual in whom the certain genomic position was covered. Following merging of these individual BAM files into one, only 7.49% of the total possible genomic positions remained without any mapped read. The distribution of these regions showed strong correlation with unassembled regions (N’s) of the genome (Fig. 2), that is, ~4.97% for GRCh38.p10 (https://www.ncbi.nlm.nih.gov/grc/human/data?asm=GRCh38.p10). The remaining uncovered positions were likely reads unmappable even to the assembled portion of the human genome, which generally consists mainly of segmental duplications, transposable elements and structural variants (Li and Freudenberg 2014). Further reduction in covered genomic positions, down to 88.27%, was the result of a filtering step that excluded from further analyses those regions having read depths below our arbitrarily set threshold of 100 mapped reads. When considering a typical human exome (Lek et al. 2016), the overall coverage of our merged data set reached 97.66% before and 94.33% after the filtering for read depth.

Variant calling was finally performed from this filtered data set, while given the number of included samples, only alleles with detected minor allele frequencies (MAF) above 5% were considered for further statistical and experimental analyses. However, we anticipate that increasing the number of aggregated samples would allow to safely decrease this threshold even to rare variants which seems to be feasible especially when considering the readily increasing worldwide popularity of NIPT testing (Green et al. 2017; Shendure et al. 2017). Besides typical SNVs, we detected also several indel variants, mainly having lengths <6bp. From these, shorter indels were found to be the most common that is in line with ExAC data reporting 95% of indels having length <6bp (Lek et al. 2016). Further comparison to large-scale studies, such as the 1000 Genomes Project (Altshuler et al. 2012; Auton et al. 2015) and the ExAC project (Lek et al. 2016), did not reveal underrepresentation of indels compared to SNVs (Fig. 3d). This suggested that one of the inherent limitations of sequencing of free-fetal DNA, stemming in the inability to detect larger indels, is not specifically relevant for NIPT-based population studies.

Since nearly 98% of the identified variants were known variants having unique entries in dbSNP, we were able to perform verification of our results on several levels. Both general and gene specific *in silico* population frequency comparisons, as well as validation analyses using a gold-standard method (Sanger sequencing) revealed high reliability of variant calling and allelic frequency determinations from our NIPT data. On the other hand, we identified 137,580 (2.08%) variants which were found to be not described in dbSNP. Frequency distributions of these novel variants were biased towards lower frequencies, when compared to dbSNP-known variants (Fig. 3c), being in line with large population studies reporting novel variants having typically low frequencies (Lek et al. 2016; van Rooij et al. 2017). Interestingly, the genomic landscape of our novel variants revealed both uniformly as well as non-uniformly distributed components (Fig. 2). Moreover, relative proportions of SNVs and indels among novel variants reflected neither proportions of previous studies (Altshuler et al. 2012; Auton et al. 2015; Lek et al. 2016), nor proportions in our dbSNP-known group, being enriched both with complex variants (51.93%) and indels (22.59%) against SNVs (25.48%)(Fig. 3d). Further analyses uncovered uniform distribution of complex variants throughout the genome (Fig. 2/Track 4), while variants identified in homopolymer and repeat-rich regions, accounting for 63.63% of novels, had non-uniform genomic distribution showing striking clustering to/near to unassembled/centromeric regions of GRCh38 (Fig. 2/Track 5). Although indel errors for Illumina platforms are considered rare in the sequencing phase itself (Nielsen et al. 2011), potential PCR amplification and realignment errors (Barnett et al. 2011; Green et al. 2017), together with low complexity genomic regions, were previously found to be typical sources of sequencing or variant calling errors covering the vast majority of false indel calls (Green et al. 2017). With this regard, given that a great reduction of overall and centromere related N’s in GRCh38 against GRCh37 stemmed in low complexity and repeat rich regions (Li and Freudenberg 2014), it cannot be considered surprising that 22.14% (30,466) of our novels failed to map back to GRCh37, pointing thus to their uniqueness to GRCh38 that, on the other hand, agreed well with previous reports (Green et al. 2017). Although this variant group may represent likely general findings, possibly consisting of both false-positives and real variants, these variants are unknown for dbSNP because of large human genome-related projects, which fuelled dbSNP, relied on variant calling against GRCh37. It should be noted, however, that none of the above characterized sources of “novelty” could be attributed to the NIPT origin of our population data and that the proportion of novel variants identified in our data set have rather technical than biological/population specific reasons.

Inherent limitations of our method should, however, also be discussed here and kept in mind during its possible future implementation. The first one is based on a recent report questioning the credibility of low-frequency variants in massively parallel sequencing based data sets, including the 1000 Genomes Project and The Cancer Genome Atlas, because of mutagenic DNA damage affecting the template DNA molecules (Chen et al. 2017). Since our paired-end reads did not overlap with each other because of short read lengths (35 bp), it is not possible to measure the extent of the described damage in our sample set. This effect could, however, be significantly reduced using DNA repair enzymes before library preparation that will most likely became a basic step in template preparations in general (Chen et al. 2017). The second possible limitation is the sample set itself. It is strongly biased towards females, since it exclusively contains women in reproductive age. It is worth noting, however, that although the original sample set contains exclusively blood samples from women, approximately 15% of reads (based on average fetal fraction), and thus also of observed alleles, belong to the fetus comprising both maternally and paternally inherited alleles. In addition, depending on the policy and possibilities of NIPT testing in each country, the sample set could be biased towards not fully physiological pregnancies, further interfering with an ideal concept of a random population sample. Since non-random population structures, based even on disease-focused consortia, are typical for large scale projects too (Lek et al. 2016), and NIPT is likely going to replace conventional screening for selected chromosomal anomalies (Gregg et al. 2016),neither of the above-mentioned concerns should be considered for absolute limitations. They should rather be considered and kept in mind when using NIPT-derived allelic frequencies in downstream applications such as in case of other large population studies.

On the other hand, large advantage of NIPT-based data lies in the fact that variants identified by low-coverage sequencing are practically not interpretable for individual patients. They become interpretable only in a statistical context when they are merged into a sufficiently large data set. Our approach, in addition, allows also de-identification of the included samples by removing all the read specific metadata from the individual files. Therefore, issues of possible genetic privacy breaching through re-identification of individual patients (Erlich and Narayanan 2014) appear to be irrelevant for NIPT derived data that simplifies the consenting phase allowing truly anonymized/pseudonymized genomic data usage for general biomedical research.

It is undisputable that large-scale reference data sets of human genetic variation are crucial for different biomedical applications. Since NIPT is globally available and the number of tests carried out rapidly increase each year (Minear et al. 2015; Gregg et al. 2016), the key important advantage of our method stems in the fact that it does not require any direct costs to generate data for large-scale population studies. It simply re-uses data already generated for other objectives thus representing a cost-effective alternative to large-scale population-specific genomic studies. Moreover, extensive cross-country data aggregation of NIPT results would represent an unprecedented source of information about worldwide frequencies of genomic variation.

## Methods

### Data source

The laboratory procedure used, to generate the NIPT data, were as follows: DNA from plasma of peripheral maternal blood was isolated for NIPT analysis from 1,548 pregnant women after obtaining a written informed consent consistent with the Helsinki declaration from the subjects. The population cohort consisted from women in reproductive age between 17-48 years with a median of 35 years. Genomic information from a sample consisted of maternal and fetal DNA fragments; 761 male, 742 female, 45 twins. Each included individual agreed to use of their genomic data in an anonymized form for general biomedical research. The NIPT study (study ID 35900_2015) was approved by the Ethical Committee of the Bratislava Self-Governing Region (Sabinovska ul.16, 820 05 Bratislava) on 30^th^ April of 2015 under the decision ID 03899_2015. Blood samples were collected to EDTA tubes and plasma was separated in dual centrifugation procedure. DNA was isolated from 700 μl of plasma using DNA Blood Mini kit (Qiagen, Hilden, DE) according to standard protocol. Sequencing libraries were prepared from each sample using TruSeq Nano kit HT (Illumina, San Diego, CA, USA) following standard protocol with omission of DNA fragmentation step. Individual barcode labelled libraries were pooled and sequenced using low-coverage whole-genome sequencing on an Illumina NextSeq500 platform (Illumina, San Diego, CA, USA) by performing paired end sequencing of 2×35 bases (Minarik et al. 2015).

### Data analysis

#### Mapping

Quality of sequenced reads was validated using reports from FastQC (v0.11.5)(Andrews 2010). Subsequently, Fastq files were mapped to human genome reference, version GRCh38.p10, using Bowtie2 (v2.1.0)(Langmead and Salzberg 2012) resulting in one SAM file for each sample. SAM files were converted to BAM format, sorted and indexed using Samtools view, merge and index utilities (v0.1.19)(Koster and Rahmann 2012). RG header with sample, lane and flow-cell identifiers was included using in-house scripts, allowing for unambiguous identification of origin of each read.

#### Exclusion of overlapping reads

Multiple observation of a single allele from the same individual may skew frequencies of variant calls. To estimate the effect, we simulated the worst-case scenario, where all overlapped reads of an individual originate from the same haplotype as may happen in case of excessive PCR duplications. Its variance was compared with more precise approach without multiple reading of a single genomic position of the same haplotype. We assigned 3 haplotypes for each individual from the 1,548 samples, corresponding to two maternal and one fetal haplotype inherited from the father. Proportions of haplotypes of an individual were determined in accordance to fetal fraction estimates from the NIPT trisomy testing (mean 14.45%, SD=2.50%). We randomly assigned reference-and alternative allele to the resulting 4,644 haplotypes, so that the proportion of the alternative ones matched the target MAF. We simulated sequencing process by gradually selecting samples and their alleles. Samples were selected randomly, without repetition, with probabilities proportional to their read count. Number of sample reads that cover the allele was selected randomly according to the coverage distribution of the sample. We considered only the most common coverages up to 3. In the worst-case simulated scenario, all selected reads covered an allele from the same haplotype. In the control scenario, we randomly picked alleles from the 3 individual’s haplotypes without repetition. Reads were generated until targeted read depth was reached. The proportion of observed alternative alleles was then recorded as simulated MAF. We repeated the simulation 1,000x for each targeted coverage and MAF. To remove all overlapping reads from individual BAM files, reads were filtered by a custom Python script in such way that only the first of overlapping reads was kept for further analysis.

#### Quality control and filtering

Summary statistics of mapping were generated using Qualimap (v2.2.1)(Okonechnikov et al. 2016). Fragments with low mapping quality (MAPQ < 21) or inconsistent mapping of corresponding reads in pair were removed using Bamtools (Barnett et al. 2011). Filtered BAM files were merged into a single BAM file using the samtools merge. Summary coverage statistics were collected with the Bedtools genomecov tool (v2.26.0)(Quinlan and Hall 2010).

#### Realignment

Positions around indels were locally realigned using RealignerTargetCreator and IndelRealigner tools from the GATK suite (DePristo et al. 2011) to minimize alignment artefacts that could lead to erroneous calls.

#### Genomic coverage

We aggregated read depth for each genomic position generated by Samtools mpileup tool (v 1.3.1) to retrieve summary coverage. We also constrained extraction to regions from the ExAC study (downloaded from ftp://ftp.broadinstitute.org/pub/ExACrelease/release0.3.1/resources/exomecallingregions.v1.intervallist) with an additional parameter – the positions. Region file has been converted into GRCh38 coordinates using Crossmap (v0.2.5)(Li et al. 2009).

#### Variant calling

Variants were identified using VarDict caller (1.5.1)(DePristo et al. 2011). We excluded variants with low allele frequency (MAF < 0.05) or low number of supporting reads (AC < 5). Called variants were checked against strand bias and converted to VCF format using teststrandBias.R and var2vcf_valid.pl script from the VarDict suite. We excluded filters for STR bias, same position in read, mean position of variants in read and strand location, because they were not suitable for our read collection of mixed population of non-overlapping, short (35 bp) reads.

#### Comparison with dbSNP

Variants were annotated against dbSNP (Sherry et al. 2001) (v150, downloaded from ftp://ftp.ncbi.nih.gov/snp/organisms/human9606b150GRCh38p7/VCF/All20170710.vcf.gz) using GATK suite. Types of individual variants (SNV, Insertion, Deletion and Complex)were inferred from TYPE attribute in the INFO field. Only variants with rs# identifier in ID field of the VCF were marked as present in dbSNP. Genomic coordinates of missing variants were converted to the GRCh37 coordinates using Crossmap. Positions that failed to map back to GRCh37 were considered as novel for the GRCh38 assembly.

#### Comparison with ExAC data

Validation of our results was performed by the comparison of selected statistical values to those extracted from the freely available ExAC data set. Simple graphical comparison of the distribution of numbers of variants with certain frequencies in the ExAC non-Finnish European population to the allelic frequencies identified in the Slovak population (Central Europe) was performed. Subsequently, a two-sample Kolmogorov-Smirnov test for testing differences between the two identified distributions was used.

Calculated frequencies for the Slovak population were compared also with 6 populations from the ExAC study (African/African American, American, East Asian, Finnish European, non-Finnish European and South Asian). Variants with at least 100 allele observations for each population from the ExAC study were extracted and compared with our frequencies determined for the Slovak population using principal component analysis implemented in Python Sklearn framework (Pedregosa et al. 2011).

### Validation of selected genomic positions using Sanger sequencing

Altogether 58 samples with good read quality were selected from our NIPT biobank. Buffy coat was separated from the deposited blood samples by centrifugation. Total DNA from buffy coat was extracted with QIAamp DNA Blood Mini Kit (Qiagen, Hilden, Germany) in compliance with the manufacturer′s instructions. Concentration of isolated DNA samples were measured on a Qubit 2.0 Fluorometer using Qubit^®^ dsDNA HS Assay Kit (Termo Fisher Scientific, Waltham, MA USA) with average DNA concentration 43ng/μl. Conventional Sanger sequencing was used to validate variant positions rs2286939(NC_000003.12:g.37020549T>C; NM_000249.3:c.1038+86T>C), rs1537514 (NC_000001.11:g.11788011G>C; NM_001010881.1:c.3812G>C), rs1800629 (NC_000006.12:g.31575254G>A; NM_000594.3:c.-488G>A), rs1801133 (NC_000001.11:g. 11796321G>A; NM_001330358.1:c.788C>T) and rs231775 (NC_000002.12:g.203867991A>G; NM_001037631.2:c.49A>G”) in 45, 12, 10, 11 and 9 DNA samples, respectively. These loci, with their surrounding regions, were PCR-amplified (primer sequences available upon request) using a HotStarTaq^®^ Master Mix Kit (Qiagen, Hilden, Germany) and the manufacturer′s protocol. Amplicon quantification was performed using Qubit 2.0 Fluorometer and Qubit dsDNA HS Assay Kit. Amplified products were verified using 2% agarose gel electrophoresis and visualised by ImageQuant LAS 500 (GE Healthcare Life Sciences). Amplicons were cleaned up by ExoSAP-IT (Thermo Fisher Scientific, Waltham, MA USA) and subsequently sequenced using a BigDye Terminator v3.1 cycle sequencing kit (Thermo Fisher Scientific, Waltham, MA USA) on Applied Biosystems ABI 3500 Genetic Analyzer.

### Data access

The sequencing reads that were used for this study are available from TrisomyTest Ltd. but restrictions apply to the availability of these data, which were used under license for the current study, and so are not publicly available. Data are however available from the authors upon reasonable request and with permission of TrisomyTest Ltd. Called variants, that support the findings of this study, are available in VCF format from https://sites.google.com/view/snipt under a flag “SNIPT”. Identified variants with corresponding information were submitted to dbSNP under the following identifiers; Handle: BIOINFKMBFNSUNIBA, Batch id: 1062867. Details of each bioinformatics step, together with the used codes and commands, are available in the *Online methods* section of this manuscript.

## Acknowledgements

This work was supported by the project titled “REVOGENE-Research Centre for Molecular Genetics” (ITMS 26240220067) supported by the Operational Programme Research and Development funded by the European Research and Developmental Fund. We would like to thank to Prof. Jozef Gecz and Dr. Mark Corbett from the University of Adelaide, Australia, for their kind help in proofreading of the manuscript and for their helpful comments. We would also like to thank to the participants allowing us to re-use their NIPT data for our project.

## Author contributions

BJ, GJ and DF designed and performed the data analyses, wrote the online methods section and proof-read the manuscript; HM, GI, SL and FR performed wet laboratory work and validation experiments based on Sanger sequencing; RJ designed the analyses, analysed the results and wrote the manuscript; GM and SM performed the routine NIPT tests as well as handled the biological material; ST proposed the leading idea of the project, designed the analyses, supervised the work and performed proofreading of the manuscript.

---

**Supplementary Figure 1:**
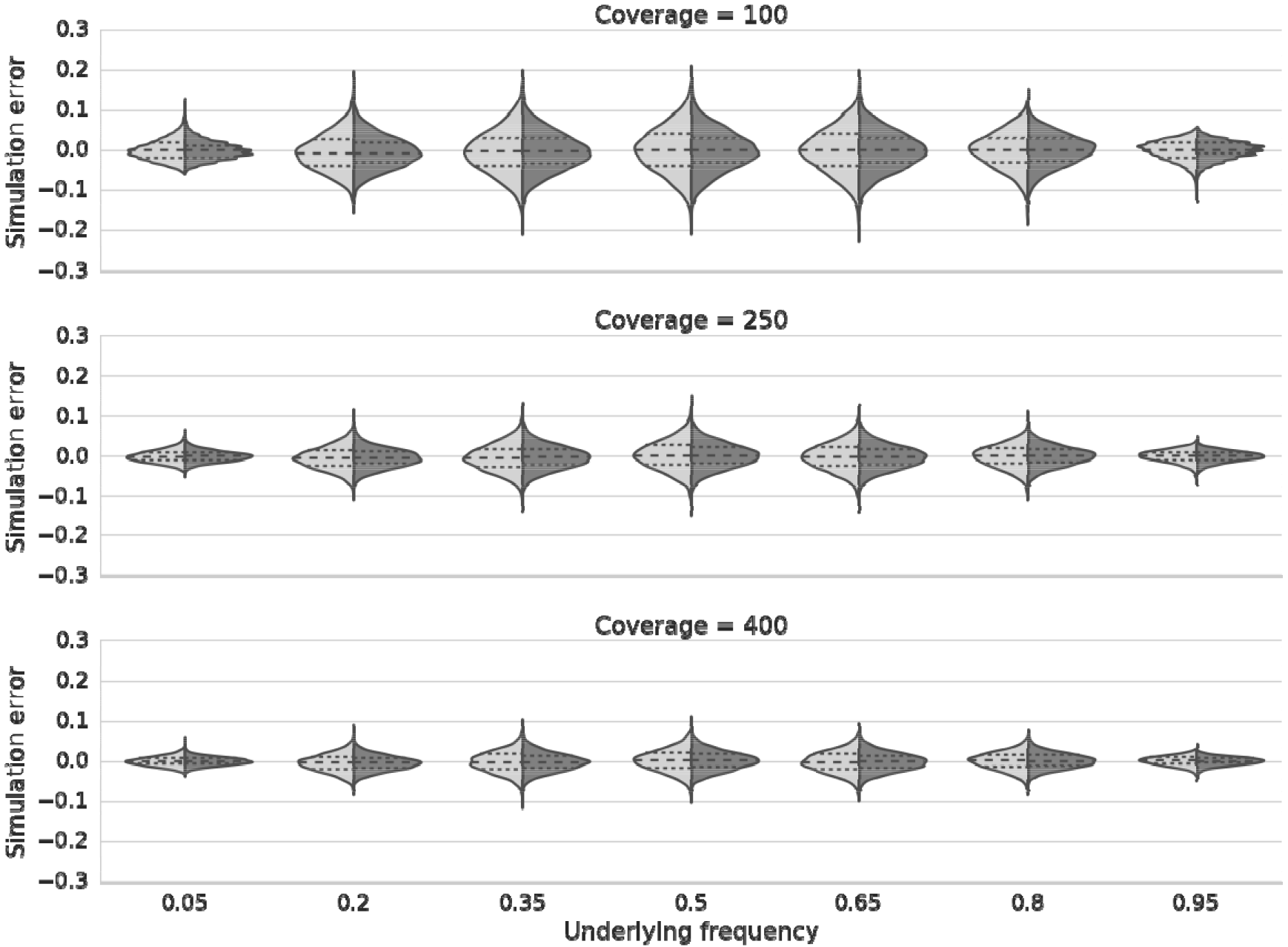
Estimated precision of allele frequency measurement from statistical simulations. Accuracy is highly dependent on the number of observed alleles and underlying allele frequency in the population. Measurements from the worst-case scenario, where all overlapped reads correspond to PCR duplicates (light grey hill) have higher dispersion than the best-case scenario where each overlapping read originates from different allele (two maternal and one fetal; dark hill).

**Supplementary Figure 2:**
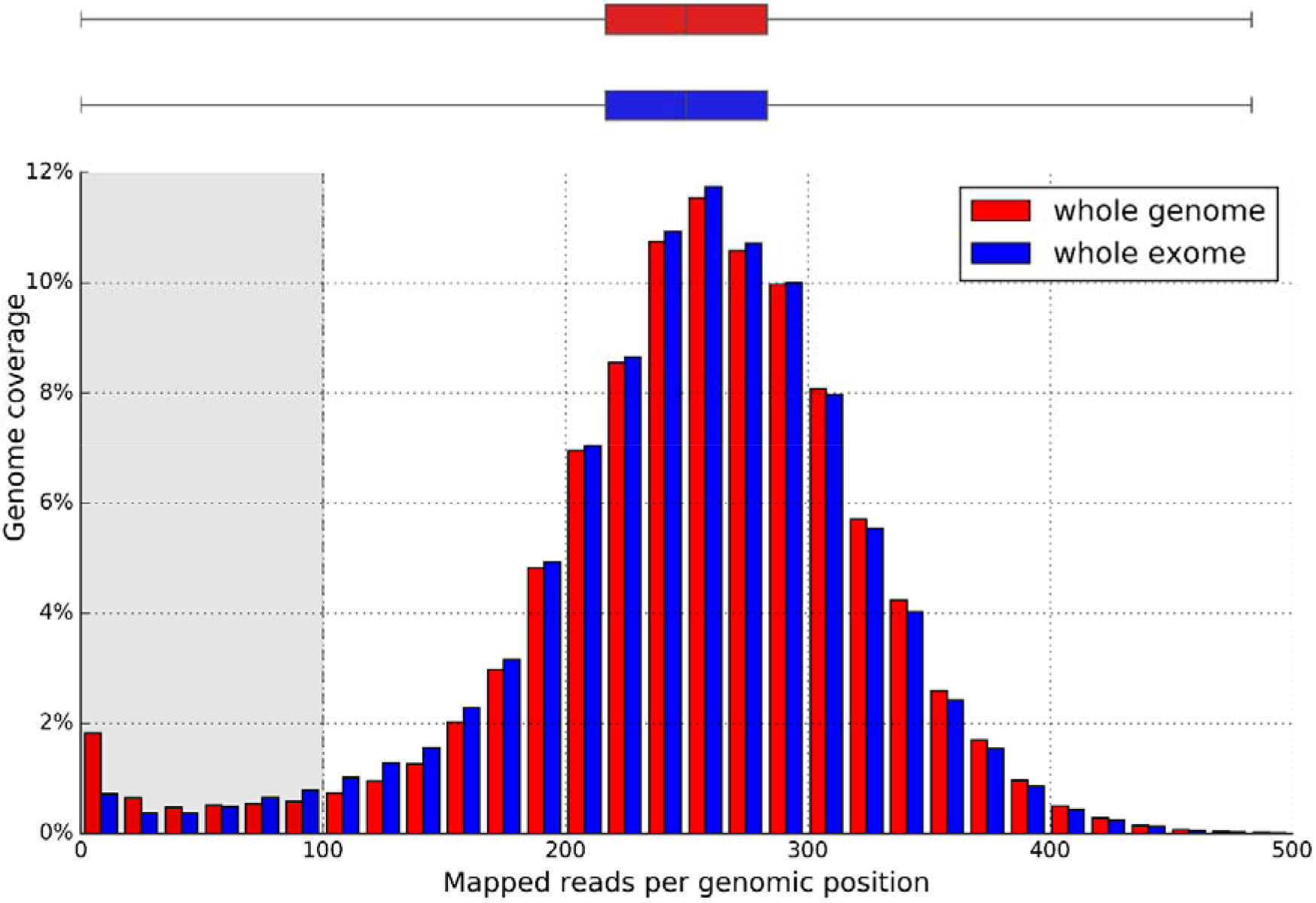
Read depth distribution with regards to the portion of the genome (exome) covered in the aggregated 1548 sample set. Positions with no mapped reads at all (7.49% for whole-genome and 2.34% for whole-exome) are not shown, while those having insufficient read depth (less than 100 reads mapped) are highlighted by the vertical grey region. Whiskers of the box plot show minimum and maximum values detected.

**Supplementary Figure 3:**
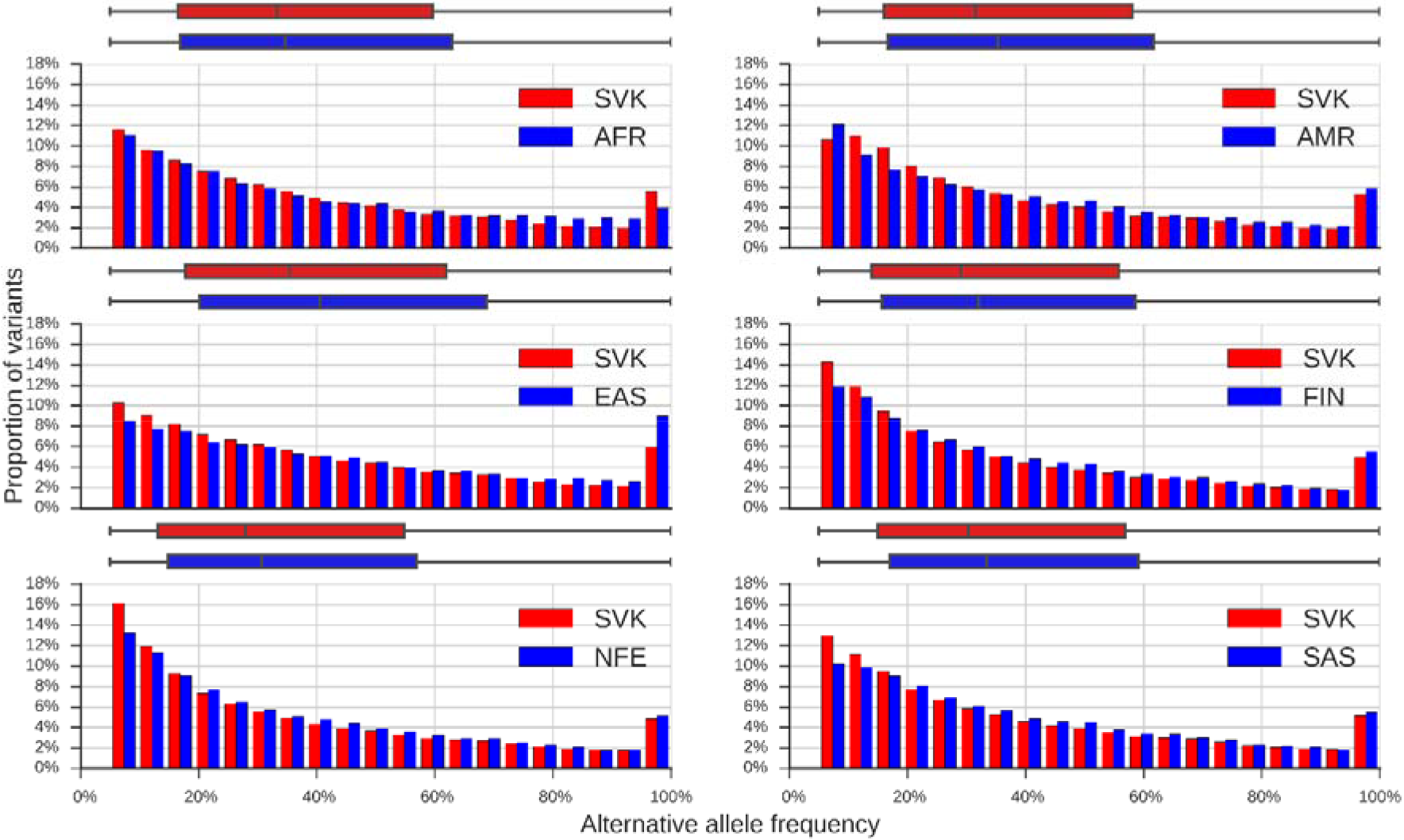
Graphical comparison between the alternative allele frequency distributions identified in our sample set and those calculated for the six ExAC population individually. Based on 71,235 variants simultaneously identified in each data subset with MAF higher than 5%. AFR = African/African American, AMR = American (Latino), EAS = East Asian, FIN = Finnish, NFE = Non-Finnish European, SAS = South Asian, SVK = Slovak.

**Supplementary Figure 4:**
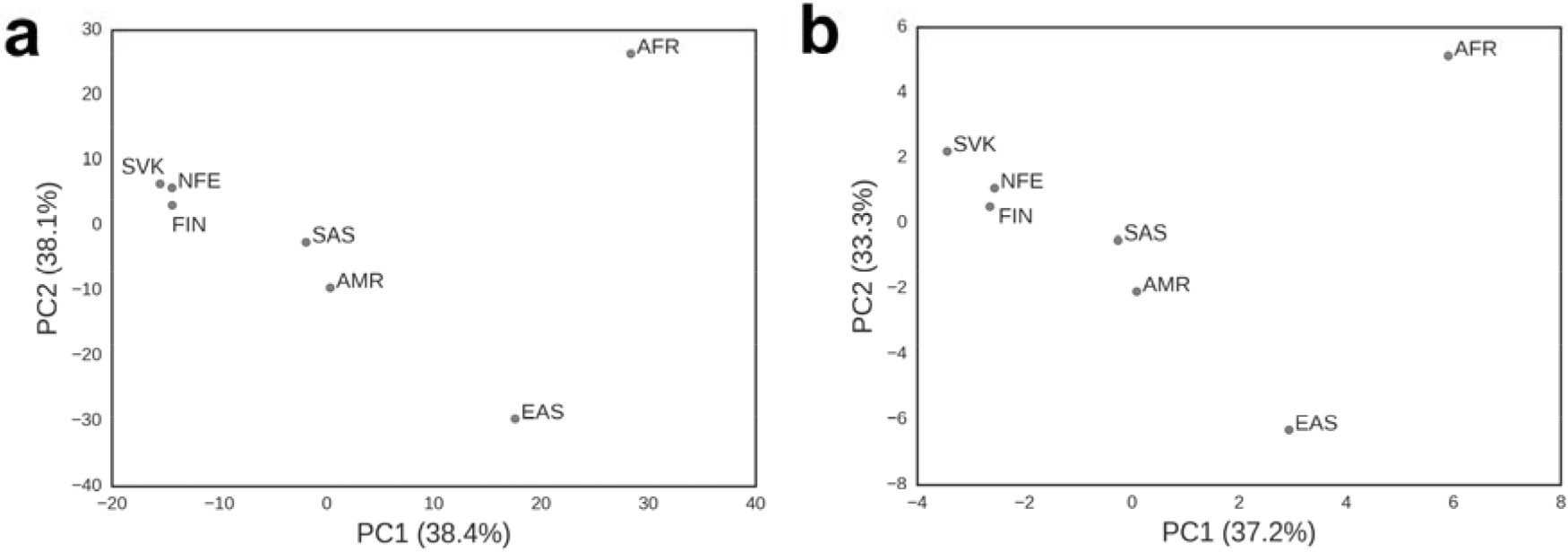
Principal component analysis (PCA) for the comparison of allelic frequencies of variants in our sample set and six different ExAC populations. Results are shown separately for (**a**) SNVs based on 68,326 variants; and (**b**) indels based on 2,909. In both cases, only those variants were used, which were simultaneously identified in each data subset with MAF higher than 5%. AFR = African/African American, AMR = American (Latino), EAS = East Asian, FIN = Finnish, NFE = Non-Finnish European, SAS = South Asian, SVK = Slovak.

**Supplementary Table 1**(uploaded as a separate Excel sheet of the Supplementary Table file): List of variants identified in the region of the CLCN1 (chloride voltage-gated channel 1, OMIM * 118425) gene in ExAC populations, dbSNP and in our data set. Only 17 of them had ExAC frequencies above 5% (highlighted in yellow). All but five were identified in our data set too with very similar calculated population frequencies. The exceptions, rs34904831, rs191902231, rs182668076, rs2280663 and rs73726622, were found to have ExAC frequencies +/−5% (depending on population). Originally three of these variants were identified in our data set too, although they were filtered out due slightly lower than 5% frequency (Suppl.Tab.1), further suggesting the feasibility of lowering our frequency restrictions. Our data contained also 89 variants missing from ExAC (highlighted in red), but found to have each of them deep intronic positions falling outside ExAC′s BED file. AFR = African/African American, AMR = American, EAS = East Asian, FIN = Finnish, NFE = Non-Finnish European, SAS = South Asian, SVK = Slovak.

**Supplementary Table 2**(uploaded as a separate Excel sheet of the Supplementary Table file): Verification of 87 positions in five polymorphic genomic loci of 58 randomly selected samples of our sample set from which genomic DNA was available to validation purposes. Reads from positions covered by multiple reads in individual samples (such in case of sample ID4730, where both alleles were identified by two different reads) were filtered out during data processing. This ensured only one allele being identified and counted from one individual in a certain position. T/C = sample originally having covered the respective position by two reads, one identifying the T allele while the other one the C allele; NIPT = non-invasive prenatal testing; -= not analysed.

